# In vitro characterization of engineered red blood cells as potent viral traps against HIV-1 and SARS-CoV-2

**DOI:** 10.1101/2020.12.20.423607

**Authors:** Magnus A. G. Hoffmann, Collin Kieffer, Pamela J. Bjorkman

## Abstract

Engineered red blood cells (RBCs) expressing viral receptors could be used therapeutically as viral traps as RBCs lack nuclei and other organelles required for viral replication. Here we show that the combination of a powerful erythroid-specific expression system and transgene codon optimization yields high expression levels of the HIV-1 receptors CD4 and CCR5, as well as a CD4-glycophorin A (CD4-GpA) fusion protein on enucleated RBCs. Engineered RBCs expressing CD4 and CCR5 were efficiently infected by HIV-1, but CD4 or CD4-GpA expression in the absence of CCR5 was sufficient to potently neutralize HIV-1 in vitro. To facilitate continuous large-scale production of engineered RBCs, we generated erythroblast cell lines stably expressing CD4-GpA or ACE2-GpA fusion proteins, which produced potent RBC viral traps against HIV-1 and SARS-CoV-2. Our results suggest that this approach warrants further investigation as a potential treatment against viral infections.

## Introduction

Red blood cells (RBCs) exhibit unique properties that can be exploited for therapeutic applications: they are the most abundant cell type, permeate all tissues, and have a lifespan of 120 days, making them attractive carriers for the delivery of therapeutic cargoes^1,2^. Moreover, RBCs do not express class I major histocompatibility complex molecules^3^, thus therapeutic RBCs from type O-negative blood could be universally administered to patients.

Engineered RBCs have been proposed as ideal candidates for the design of viral traps, as they lack nuclei and other organelles required for viral replication^4-7^. Viruses could be lured into attaching to and infecting RBCs that present viral receptors, thereby leading the virus to a dead end and protecting viral target cells from infection. This approach has the potential to prevent viral escape, as viruses must retain the ability to bind their receptors. However, expression of viral receptors on RBCs is difficult to achieve since mature erythrocytes lack the cellular machinery to synthesize proteins. Hence erythroid progenitor cells need to be genetically-engineered to express the viral receptors and then be differentiated into enucleated RBCs. During the erythroid differentiation process, transgene expression is restricted through transcriptional silencing^8^, translational control mechanisms^9^ and degradation of proteins that are not normally present in RBCs^10^.

Here we show that the combination of a powerful erythroid-specific expression system and transgene codon optimization yields high expression levels of the HIV-1 receptors CD4 and CCR5 on enucleated RBCs to generate viral traps that potently inhibit HIV-1 infection in vitro. We then applied these engineering strategies to generate erythroblast cell lines that can continuously produce potent RBC viral traps against HIV-1 and SARS-CoV-2.

## Results

### Enucleated RBCs express HIV-1 receptors

We used an in vitro differentiation protocol^11^ to differentiate human CD34+ hematopoietic stem cells (HSCs) into reticulocytes, an immature form of enucleated RBC that still contains ribosomal RNA (Fig. 1a). At the end of the proliferation phase, erythroid progenitor cells were transduced using lentiviral vectors carrying CD4 or CCR5 transgenes by spinoculation (Fig. 1a; Supplementary Fig. 1a). We also evaluated expression of a CD4-glycophorin A (CD4-GpA) fusion protein that contained the extracellular CD4 D1D2 domains fused to the N-terminus of GpA, an abundantly-expressed RBC protein. This strategy enabled expression of single-domain antibodies in RBCs^11^, but is applicable only to CD4, a single-pass transmembrane protein, but not to CCR5, a multi-pass transmembrane protein. Three days post-transduction, transgene expression was evaluated by flow cytometry. Expression was low for transgenes when the CMV promoter or alternative ubiquitous promoters were used (Fig. 1b; Supplementary Fig. 1b). CD4-GpA expressed only marginally better than CD4, suggesting that additional strategies are required to achieve robust expression of viral receptors.

**Fig. 1.**
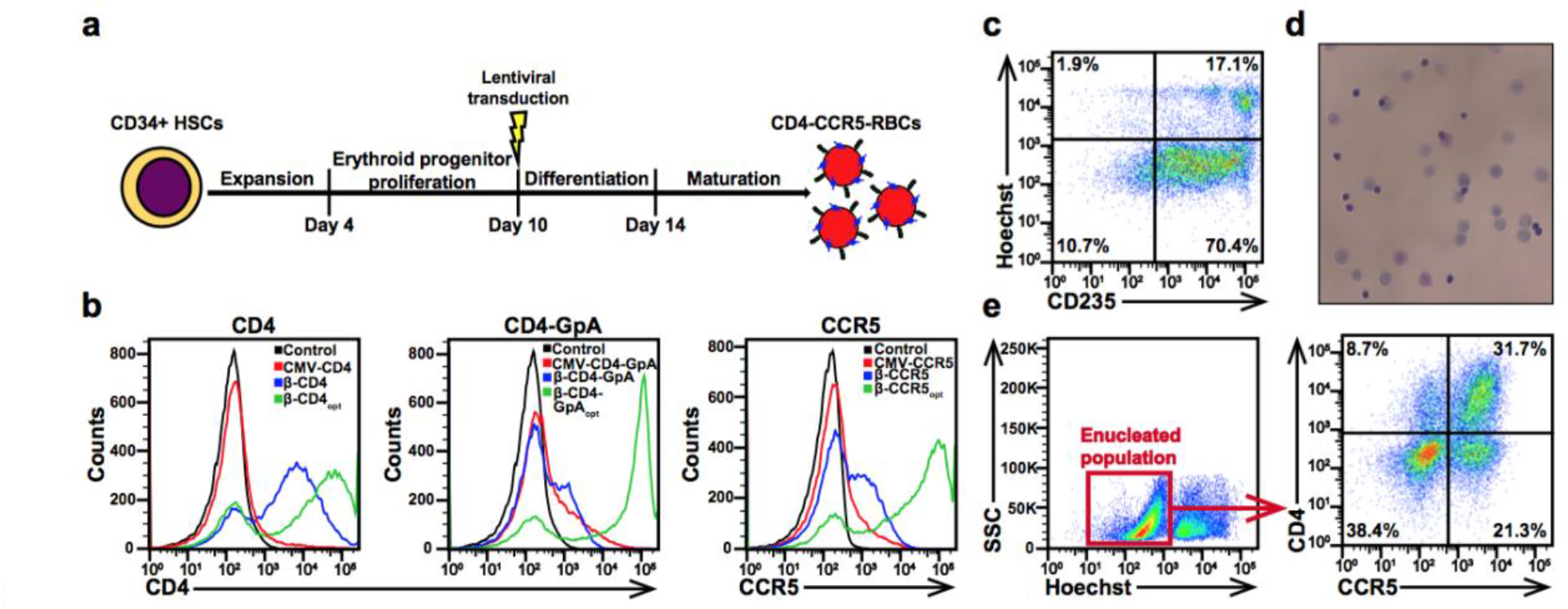
Engineered RBCs express HIV-1 receptors. **a**, Schematic illustrating the workflow for generating enucleated RBCs expressing HIV-1 receptors. **b**, Flow cytometry analysis of CD4, CD4-GpA, and CCR5 expression on day 13 of differentiation comparing the CMV promoter (red), the β-globin promoter (blue), and the β-globin promoter in combination with codon optimization (green). **c**, Quantification of enucleated CD4-CCR5-RBCs by flow cytometry. Enucleated RBCs expressed CD235 and did not stain for the nuclear dye Hoechst. **d**, Image of CD4-CCR5-RBCs after May-Grunwald-Giemsa staining (magnification x63). **e**, CD4 and CCR5 expression on enucleated (Hoechst-negative) RBCs.

To evaluate whether transcriptional silencing can be prevented by using an erythroid-specific promoter, transgenes were subcloned into the CCL-βAS3-FB lentiviral vector^12^, which contains regulatory elements that support the high expression levels of β-globin during erythroid development (vectors β-CD4, β-CD4-GpA, and β-CCR5) (Supplementary Fig. 1a). CD4 expression was greatly enhanced by this expression system, CCR5 expression increased to a lesser extent, but CD4-GPA expression was not improved (Fig. 1b). We hypothesized that the limited availability of ribosomes and transfer RNAs restricted transgene expression in differentiating erythroid cells and therefore codon-optimized transgene cDNA sequences (β-CD4_opt_, β-CD4-GpA_opt_, and β-CCR5_opt_), resulting in greatly enhanced expression for all transgenes (Fig. 1b).

Genetically-engineered CD4+/CCR5+ erythroid progenitor cells differentiated efficiently into enucleated RBCs (Fig. 1c). After differentiation, almost all cells expressed GpA, of which >80% did not stain for Hoechst nuclear dye, and May-Grunwald-Giemsa staining confirmed that the majority of cells were enucleated RBCs (Fig. 1c,d). Approximately 1/3 of enucleated RBCs expressed CD4 and CCR5 (Fig. 1e) at levels comparable to a CD4+ T-cell line (Supplementary Fig. 2).

### Engineered RBCs efficiently entrap and neutralize HIV-1

To evaluate the efficacy of RBC viral traps against HIV-1, we generated RBCs that expressed CD4+/-CCR5 or CD4-GpA+/-CCR5 (Fig. 2a) and used the β-lactamase (BlaM) fusion assay^13^ to evaluate if HIV-1 can infect RBC viral traps. RBCs were incubated with a CCR5-tropic HIV-1_YU2_ pseudovirus carrying a BlaM-Vpr fusion protein that enters cells upon infection. When infected cells are exposed to the fluorescence resonance energy transfer (FRET) substrate CCF2-AM, BlaM cleaves the β-lactam ring in CCF2-AM resulting in a shift of its emission spectrum from green (520 nm) to blue (447 nm)^13^. Whereas infectious events were ≤0.3% in control RBCs and CD4-RBCs, 6.1% of CD4-CCR5-RBCs were infected (Fig. 2b). Since only 1/3 of these RBCs expressed both receptors (Fig. 1e), this corresponds to infection of almost 20% of CD4-CCR5-RBCs. Higher infection rates were observed for RBCs that co-expressed CD4 and CXCR4 after incubation with a CXCR4-tropic HIV-1 HxBc2 pseudovirus (Supplementary Fig. 3a,b). However, RBCs that co-expressed the CD4-GpA fusion protein and CCR5 or CXCR4 were infected at lower frequencies (Fig. 2b; Supplementary Fig. 3b). Addition of the CD4 D3D4 domains to CD4-GpA did not improve infection rates (Supplementary Fig. 3c). Unlike CD4, GpA does not localize to lipid raft subdomains^14^, thus we speculate that low infection resulted from a lack of co-localization between CD4-GpA and the CCR5 and CXCR4 co-receptors.

**Fig. 2.**
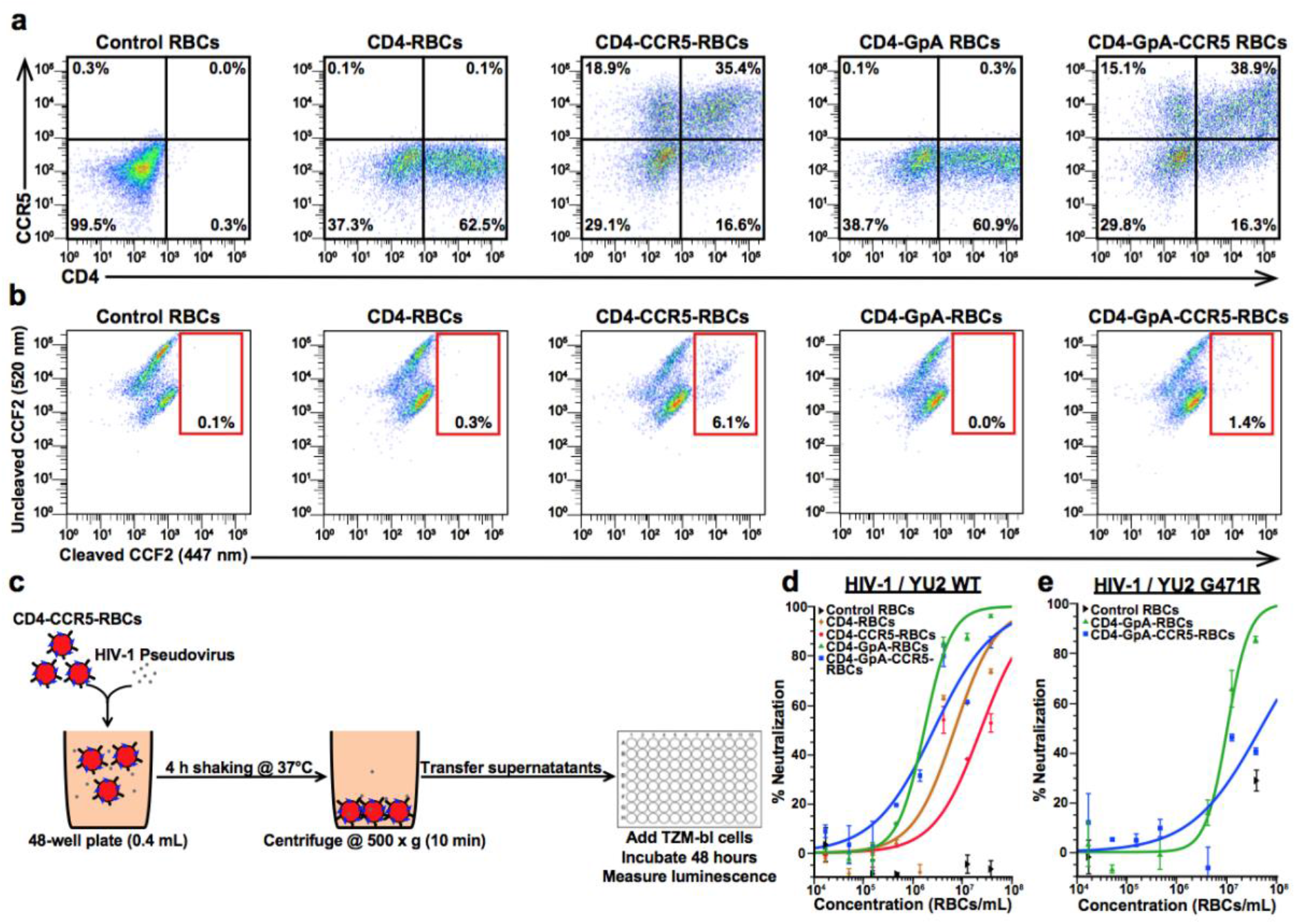
Engineered RBCs can be infected by HIV-1 and potently neutralize the virus in vitro. **a**, Flow cytometry measurement of CD4 and CCR5 expression at the end of differentiation for control RBCs, CD4-RBCs, CD4-CCR5-RBCs, CD4-GpA-RBCs, and CD4-GpA-CCR5-RBCs. **b**, Flow cytometry analysis of HIV-1 infection of engineered RBCs after overnight incubation with a HIV-1_YU2_ pseudovirus carrying a Vpr-BlaM fusion protein. BlaM cleaves the FRET substrate CCF2-AM in infected cells resulting in a shift of its emission spectrum from green (520 nm) to blue (447 nm). **c**, Schematic illustrating the workflow for the modified neutralization assay used to evaluate the neutralization activity of engineered RBC viral traps. **d**, In vitro neutralization assay against HIV-1_YU2_ pseudovirus comparing control RBCs (black), CD4-RBCs (brown), CD4-CCR5-RBCs (red), CD4-GpA-RBCs (green), and CD4-GpA-CCR5-RBCs (blue). Data points are the mean and SD of duplicate measurements. **e**, In vitro neutralization assay against mutant HIV-1_YU2_ Env G471R pseudovirus comparing control RBCs (black), CD4-GpA-RBCs (green), and CD4-GpA-CCR5-RBCs (blue). Data points are the mean and SD of duplicate measurements.

**Fig. 3.**
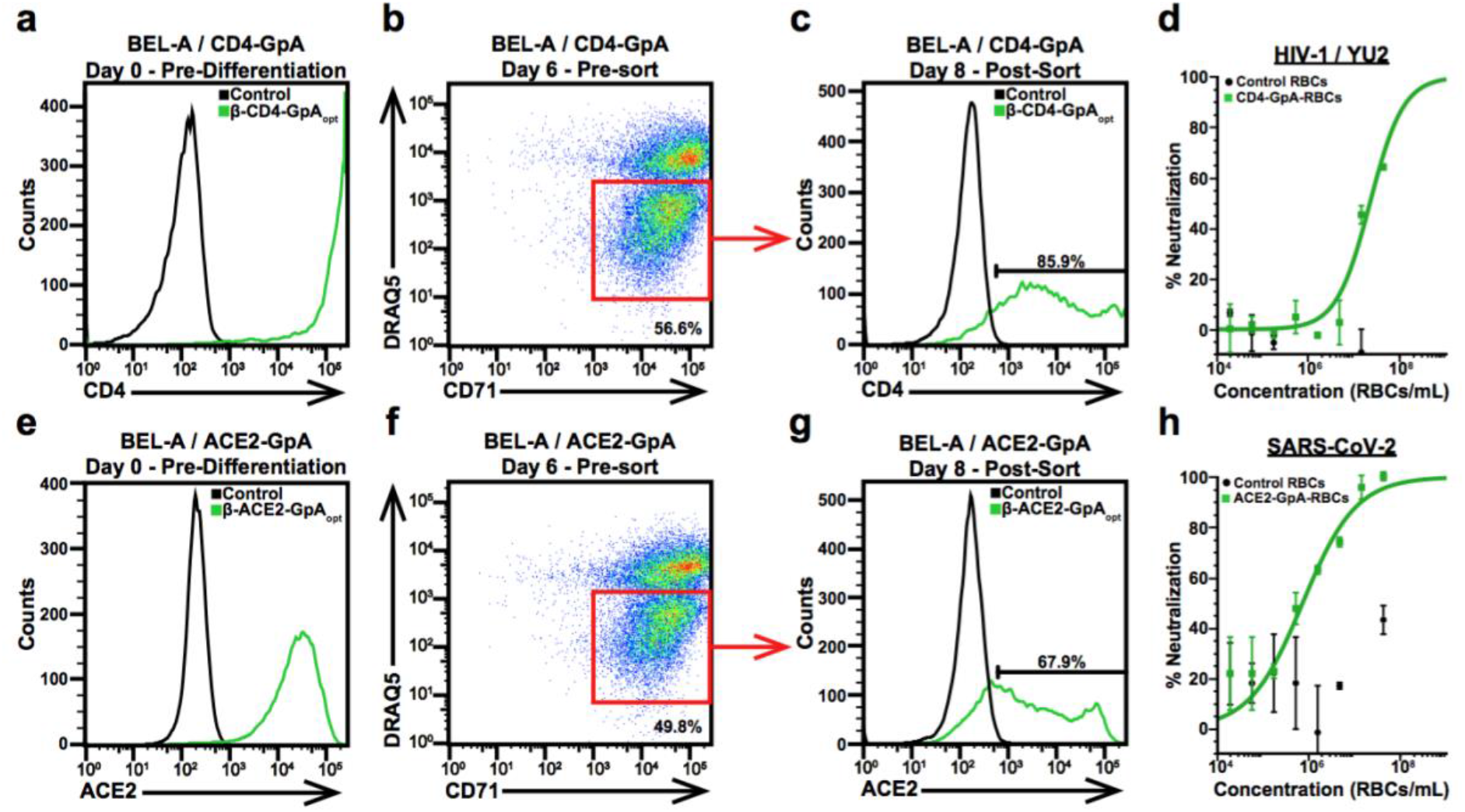
BEL-A erythroblast cell lines stably express CD4-GpA and ACE2-GpA to produce potent RBC viral traps against HIV-1 and SARS-CoV-2. **a**, Flow cytometry measurement of CD4-GpA expression on BEL-A / CD4-GpA cells pre-differentiation. **b**, Flow cytometry analysis of enucleated CD4-GpA-RBCs on day 6 of differentiation. Enucleated RBCs expressed CD71 and did not stain for the nuclear dye DRAQ5. **c**, Flow cytometry analysis of CD4-GpA expression on CD71+/DRAQ5-BEL-A / CD4-GpA cells post-sorting on day 8 of differentiation. **d**, In vitro neutralization assay against HIV-1_YU2_ pseudovirus comparing control RBCs (black) and CD4-GpA-RBCs (green). Data points are the mean and SD of duplicate measurements. **e**, Flow cytometry analysis of ACE2-GpA expression on BEL-A / ACE2-GpA cells pre-differentiation. **f**, Flow cytometry analysis of enucleated ACE2-GpA-RBCs on day 6 of differentiation. Enucleated RBCs expressed CD71 and did not stain for the nuclear dye DRAQ5. **g**, Flow cytometry measurement of ACE2-GpA expression on CD71+/DRAQ5-BEL-A / ACE2-GpA cells post-sorting on day 8 of differentiation. **h**, In vitro neutralization assay against lentivirus-based SARS-CoV-2 pseudovirus comparing control RBCs (black) and ACE2-GpA-RBCs (green). Data points are the mean and SD of duplicate measurements.

We assessed the therapeutic potential of RBC viral traps using a modified version of the HIV-1 TZM-bl neutralization assay^15^ (Fig. 2c). After incubating RBCs with HIV-1_YU2_ pseudovirus, samples were centrifuged to remove RBCs and virions that attached to or infected RBCs. Supernatants containing free virions that had not been captured by RBCs were transferred to 96-well plates and TZM-bl cells were added to measure infectivity. CD4-GpA-RBCs neutralized HIV-1_YU2_ most potently at a half-maximal inhibitory concentration (IC_50_) of 1.7×10^6^ RBCs/mL (Fig. 2d). This concentration is equivalent to 0.03% of the RBC concentration of human blood (∼5×10^9^ RBCs/mL), suggesting that it is feasible to achieve therapeutic concentrations in vivo. CD4-GpA-RBCs were ∼4-fold more potent than CD4-RBCs, likely due to higher expression levels (Fig. 2a; Supplementary Fig. 4). Surprisingly, co-expression of CCR5 lowered the neutralization activity of CD4-GpA-CCR5-RBCs and CD4-CCR5-RBCs by ∼2-3-fold, respectively (Fig. 2d), implying that HIV-1 infection of RBC viral traps is not required for potent neutralization. CCR5 co-expression slightly lowered CD4-GpA and CD4 expression levels (Fig. 2a), potentially explaining the observed drop in potency. However, these results do not exclude the possibility that CCR5 expression on RBC viral traps has beneficial effects in vivo.

We previously showed that virus-like nanoparticles presenting clusters of CD4 (CD4-VLPs) that formed high-avidity interactions with trimeric HIV-1 Env spikes on virions potently neutralized a diverse panel of HIV-1 strains and prevented viral escape in vitro^16^. To confirm that RBC viral traps can also form high-avidity interactions with Env, we evaluated neutralization against a mutant HIV-1_YU2_ Env G471R pseudovirus that was resistant to monomeric soluble CD4, but was sensitive to CD4-VLPs^16^. CD4-GpA-RBCs potently neutralized the HIV-1_YU2_ G471R pseudovirus (IC_50_=1.0×10^7^ RBCs/mL) (Fig. 2e), suggesting that RBC viral traps and CD4-VLPs would be similarly effective in preventing viral escape through formation of high-avidity interactions with HIV-1 Env spikes.

### Erythroblast cell lines stably express viral receptors and continuously produce RBC viral traps against HIV-1 and SARS-CoV-2

To generate a renewable source of RBC viral traps, we engineered the immortalized BEL-A erythroblast cell line^17^ to stably express high levels of CD4-GpA (Fig. 3a). The CD4-GpA– BEL-A cells efficiently differentiated into enucleated RBCs, as >50% of CD71-expressing cells did not stain for the nuclear marker DRAQ5 (Fig. 3b). After differentiation, CD71+/DRAQ5-RBCs were purified using fluorescence-activated cell sorting (FACS). The majority of RBCs still expressed CD4-GpA (Fig. 3c) and potently neutralized HIV-1_YU2_ (IC_50_=2.1×10^7^ RBCs/mL) (Fig. 3d).

To evaluate if RBC viral traps could be effective against other viruses, we generated a BEL-A cell line that continuously produces RBC viral traps against SARS-CoV-2, the virus that caused the ongoing COVID-19 pandemic^18^. BEL-A cells were transduced to stably express a chimeric ACE2-GpA protein containing the extracellular domain of the SARS-CoV-2 receptor ACE2^18^ fused to GpA (Fig. 3e). Differentiation efficiency and transgene expression on sorted CD71+/DRAQ5-RBCs was comparable to the CD4-GpA cell line (Fig. 3f,g). Importantly, lentivirus-based SARS-CoV-2 pseudovirus^19^ was highly susceptible to ACE2-GpA-RBC neutralization (IC_50_=7⨯ 10^5^ RBCs/mL) (Fig. 3h) suggesting that RBC viral traps have the potential to become powerful anti-viral agents against a range of viruses.

## Discussion

In summary, we described engineering strategies that facilitate efficient and continuous production of potent RBC viral traps against HIV-1 and SARS-CoV-2. Engineered RBCs expressing HIV-1 receptors were efficiently infected by HIV-1 and potently neutralized the virus in vitro, thus demonstrating the desired properties of a viral trap. Expression of CD4 or the CD4-GpA fusion protein in the absence of CCR5 was sufficient to potently neutralize HIV-1 due to the formation of high-avidity interactions between clusters of CD4 or CD4-GpA on the RBC surface and trimeric HIV-1 Env spikes on virions. We have previously shown that such high-avidity interactions enhanced the potency of CD4-VLPs by >10,000-fold in comparison to conventional CD4-based inhibitors such as soluble CD4 and CD4-Ig, and HIV-1 was unable to escape against CD4-VLPs in vitro^16^. In contrast to CD4-VLPs, RBC viral traps would persist in vivo for months, implying the RBC approach has the potential to provide sustained control of HIV-1 infection.

Engineered RBCs expressing the ACE2-GpA fusion protein potently neutralized SARS-CoV-2 suggesting that RBC viral traps could be an effective treatment against a diverse range of viruses. Since cell lines that generate RBC viral traps could be rapidly developed once a host receptor has been identified, RBC viral traps could also become a rapid-response treatment strategy for future viral outbreaks. Our results suggest that this approach warrants further investigation as a potential treatment against viral infections.

## Methods

### In vitro CD34+ HSC differentiation

Human cord blood or mobilized peripheral blood CD34+ HSCs (StemCell Technologies) were differentiated into enucleated RBCs using a modified version of a previously-described protocol^11^. Briefly, CD34+ HSCs were cultured in expansion medium (100 ng/mL rhFlt3, 100 ng/mL rhSCF, 20 ng/mL rhIL-6, 20 ng/mL rhIL-3, and 100 nM dexamethasone in StemSpan II medium) at a density of 10^5^ cells/mL for 4 days. Cells were then placed in differentiation 1-2 medium (2% human AB plasma, 3% human AB serum, 3 U/mL heparin, 10 ng/mL rhSCF, 1 ng/mL rhIL-3, and 3 U/mL erythropoietin in StemSpan II medium) at a density of 10^5^ cells/mL for 3 days and at 2⨯ 10^5^ cells/mL for an additional 3 days. The cells were then passaged into differentiation 3 medium (2% human AB plasma, 3% human AB serum, 3 U/mL heparin, 10 ng/mL rhSCF, and 1 U/mL erythropoietin in StemSpan II medium) at a density of 2 ⨯ 10^5^ cells/mL for 4 days. To induce RBC maturation, cells were cultured in differentiation 4 medium (2% human AB plasma, 3% human AB serum, 3 U/mL heparin, 0.1 U/mL erythropoietin, and 200 μg/mL holo-transferrin in StemSpan II medium) at a density of 10^6^ cells/mL for 4 days, and in differentiation 5 medium (2% human AB plasma, 3% human AB serum, 3 U/mL heparin, and 200 μg/mL holo-transferrin in StemSpan II medium) at a density of 5 × 10^6^ cells/mL for an additional 3 days. For morphological analysis, cells were spun onto glass slides by cytocentrifugation, stained with May-Grünwald-Giemsa reagents (Sigma-Aldrich), and examined under an LSM800 laser scanning confocal microscope (Zeiss).

### Transgenes and codon optimization

Human CD4, CCR5, CXCR4, ACE2 and glycophorin A (GpA) cDNA sequences were obtained from the National Center for Biotechnology Information. The CD4-GpA fusion construct encoded the CD4 signal peptide and D1D2 domains fused to the N-terminus of GpA with a 9-residue linker (Glu-Pro-Lys-Thr-Pro-Lys-Pro-Gln-Pro). The ACE2-GpA fusion protein construct encoded the extracellular domain of human ACE2 (residues 1-614) fused to the N-terminus of GpA with the 9-residue linker. Transgenes were cloned into the lentiviral backbone plasmids pHAGE-IRES-ZsGreen (PlasmID Repository, Harvard Medical School) for expression under ubiquitous promoters (CMV, EF1α, UBC, and CASI promoters) and pCCL-FB^12^ (provided by Dr. Donald Kohn, UCLA) for erythroid-specific expression. Codon optimization of transgene cDNA sequences was performed using the GeneArt GeneOptimizer software (Thermo Fisher Scientific).

### Lentiviral transduction

VSV-G-pseudotyped lentiviral vectors were produced by co-transfecting HEK293T cells with lentiviral backbone plasmids and packaging plasmids (pHDM-Hgpm2, pHDM-tat1b, pRC/CMV-rev1b, pHDM-G) using Fugene HD (Promega) according to the manufacturer’s protocol. Supernatants were collected after 48 and 72 hours, and lentiviral vectors were concentrated 50-fold using Lenti-X concentrator solution (Takara) according to the manufacturer’s protocol. On day 10 of the differentiation protocol, erythroid progenitor cells were seeded at a density of 10^6^ cells/mL in 12-well plates in the presence of 10 μg/mL polybrene. 20 μL of concentrated lentiviral vector was added per well and plates were spun for 1.5 hours at 850 × g at 30°C. Plates were then incubated for 3 hours at 37°C before passaging the transduced cells into differentiation 3 medium. For cells that were co-transduced to express two transgenes, 20 μL of each lentiviral vector was added per well. To generate large numbers of engineered RBCs for neutralization assays, two transductions steps were performed on days 10 and 14 of the differentiation protocol.

### Flow cytometry

Transgene expression and RBC maturation efficiency were analyzed by flow cytometry (MACSQuant, Miltenyi Biotec). 2-3 ⨯ 10^5^ cells were collected for each condition and samples were stained with the following antibodies: APC-conjugated anti-human CD4 (Invitrogen), FITC-conjugated anti-human CD4 (BD Bioscience), FITC-conjugated anti-human CCR5 (BioLegend), PE-conjugated anti-human CXCR4 (Invitrogen), FITC-conjugated anti-human ACE2 (R&D Systems), APC-conjugated anti-CD235ab (BioLegend), and Brilliant Violet 421-conjugated anti-human CD71 (BioLegend). The percentage of enucleated RBCs was measured by double staining cells with APC-conjugated anti-CD235ab and the nuclear stain Hoechst (Thermo Fisher Scientific). Enucleated RBCs were defined as CD235ab+/Hoechst-cells. The percentage of enucleated RBCs that expressed transgenes was measured by triple-staining cells with APC-conjugated anti-human CD4, FITC-conjugated anti-human CCR5, and Hoechst nuclear stain.

### β-lactamase fusion assay

The ability of engineered RBCs to be infected by HIV-1 was evaluated using a modified version of the β-lactamase (BlaM) assay^13^. R5-tropic HIV-1_YU2_ and X4-tropic HIV-1_HxBc2_ pseudovirus were produced by co-transfecting a confluent T75 flask of HEK293T cells with the PSG3ΔEnv backbone plasmid (8 μg), the YU2 or HxBc2 Env expression plasmid (4 μg), and a plasmid expressing a BlaM-Vpr fusion protein^20^ (4 μg; provided by Dr. Wesley Sundquist, University of Utah). The supernatant was collected after 72 hours and concentrated by centrifugal filtration. 5 ⨯ 10^4^ RBCs were seeded in 100 μL differentiation 5 medium in 96-well plates in the presence of 10 μg/mL polybrene. 20 μL of concentrated YU2-BlaM-Vpr or HxBc2-BlaM-Vpr pseudovirus were added and plates were spun at 1,000 × g for 1 hour at 30°C. Plates were then incubated at 37°C overnight. On the next day, freshly prepared 6X CCF2-AM labeling solution was added, and cells were stained for 2 hours at room temperature in the dark. After two washes with PBS, the cells were analyzed by flow cytometry (MACSQuant, Miltenyi Biotec).

### HIV-1 neutralization assays

The ability of engineered RBCs to inhibit HIV-1 infection of target cells was tested by using a modified version of the HIV-1 pseudovirus-based TZM-bl assay^15^. Briefly, serial dilutions of control and engineered RBCs were seeded in 400 μL TZM-bl media in 48-well plates and incubated with 0.4 μL HIV-1_YU2_ pseudovirus (TCID_50_ = 3.2 ⨯ 10^5^ IU/mL) for 4 hours on an orbital shaker (400 rpm) at 37°C in the presence of 10 μg/mL of polybrene. Cells were then spun down at 500 ⨯ g for 10 min and 155 μL of the supernatants were transferred to 96-well plates. TZM-bl reporter cells (NIH AIDS Reagents Program) were added, and luminescence was measured after 48 hours.

### SARS-CoV-2 neutralization assays

Lentivirus-based SARS-CoV-2 pseudovirus was generated by transfecting HEK293T cells with a luciferase-expressing lentiviral backbone plasmid, accessory plasmids (pHDM-Hgpm2, pHDM-tat1b, pRC/CMV-rev1b), and a plasmid encoding the SARS-CoV-2 Spike protein with a 21-residue cytoplasmic tail deletion (Wuhan Hu-1 strain; GenBank NC_045512). The neutralization activity of ACE2-GpA RBCs was measured using a modified version of a recently-reported protocol^19^. 1.25 ⨯ 10^4^ 293T-ACE2 cells (provided by Dr. Jesse Bloom, Fred Hutchinson Cancer Research Center) were seeded per well on poly-L-Lysine-coated 96-well plates (Corning) 18 hours before infection. Serial dilutions of control and ACE2-GpA-RBCs were seeded in 400 μL media (DMEM supplemented with 10% FBS and Pen-Strep) and incubated with 3 μL of lentiviral particles pseudotyped with the SARS-CoV-2 Spike protein (5. ⨯ 10^7^ RLU/mL) for 4 hours on an orbital shaker (400 rpm) at 37°C in the presence of 10 μg/mL of polybrene. The lentiviral backbone of this SARS-CoV-2 pseudovirus system expresses luciferase to enable detection of infected cells. RBCs were spun down at 500 × g for 10 min and 100 μL supernatant was transferred to the 96-well plate with the seeded 293T-ACE2 cells. Luminescence was measured after 48 hours using a plate reader (Tecan).

### Generation of stable erythroblast cell lines

Immortalized BEL-A erythroblast cells^17^ (provided by Dr. Jan Frayne, University of Bristol) were transduced with VSV-G–pseudotyped lentiviral vectors carrying the CD4-GpA or ACE2-GpA transgenes in the erythroid-specific pCCL-FB expression cassette. To allow positive selection of cells that stably expressed the transgenes, the puromycin-N-acetyltransferase gene was added downstream of the transgene and a P2A cleavage peptide^21^. Stable BEL-A / CD4-GpA and BEL-A / ACE2-GpA cell lines were generated by growing the transduced cells in expansion media (50 ng/mL rhSCF, 3 U/mL erythropoietin, 1 µM dexamethasone, and 1 µg/mL doxycycline in StemSpan II medium) in the presence of 0.25 µg/mL puromycin for 3-4 weeks. Differentiation of BEL-A cells was initiated as described^17^ by transferring the cells into primary media (3% human AB serum, 2% FBS, 3 U/mL heparin, 10 ng/mL rhSCF, 1 ng/mL rhIL-3, 3 U/mL erythropoietin, 200 µg/mL holo-transferrin, and 1 µg/mL doxycycline in StemSpan II medium) for 3-4 days at a density of 2 ⨯ 10^5^ cells/mL. To induce RBC maturation, cells were moved into tertiary media (3% human AB serum, 2% FBS, 3 U/mL heparin, 3 U/mL erythropoietin, 500 µg/mL holo-transferrin, and 1 U/mL Pen-Strep in StemSpan II medium) for 4 days at a density of 1 ⨯ 10^6^ cells/mL.

### Fluorescence-activated cell sorting (FACS)

Enucleated RBCs were purified by FACS on day 7 of the BEL-A differentiation protocol. Brilliant Violet 421-conjugated anti-human CD71 antibody (BioLegend) and the nuclear stain DRAQ5 (Abcam) were diluted 1:100 and 1:1,000 in PBS+ (PBS supplemented with 2% FBS). Cells were stained at a concentration of 2.5 ⨯ 10^7^ cells/mL for 30 min at room temperature in the dark. After two washes in PBS+, cells were resuspended in PBS+ at a concentration of 1 × 10^7^ cells/mL. Enucleated RBCs were defined as CD71+/DRAQ5-cells and this cell population was purified using a SONY SH800 cell sorter (Sony Biotechnology).

## Acknowledgements

We thank N.J. Huang, N. Pishesha, and H.F. Lodish for helpful discussion, advice, and reagents; P.N.P., Gnanapragasam and L.M. Kakutani for producing SARS-CoV-2 pseudovirus and setting up the SARS-CoV-2 pseudovirus neutralization assay in our laboratory; G.L. Chadwick, R. Galimidi, and A. Moradian (formerly Caltech Proteome Exploration Laboratory) for helpful discussions and reagents; G. Spigolon for guidance with light microscopy performed at the Beckman Institute Biological Imaging Facility; Z. Romero-Garcia and D.B. Kohn for the pCCL-AS3-FB plasmid; J. Frayne for BEL-A cells; J. Voetteler and W.I. Sundquist for the CCF2-AM reagent; J.D. Bloom for 293T-ACE2 cells and plasmids for generating SARS-CoV-2 pseudovirus; and the NIH AIDS Reagent Program for reagents. This work was supported by the Bill and Melinda Gates Foundation grant OPP1202246, the DeLogi Trust (facilitated by Caltech), and by a generous gift from Kairos Ventures (facilitated by Caltech).

## Author Contributions

M.A.G.H. and P.J.B. designed the research, M.A.G.H. and C.K. performed the research, M.A.G.H. and P.J.B. analyzed data, and M.A.G.H. and P.J.B. wrote the manuscript.

## Supplementary information

**Fig. S1.**
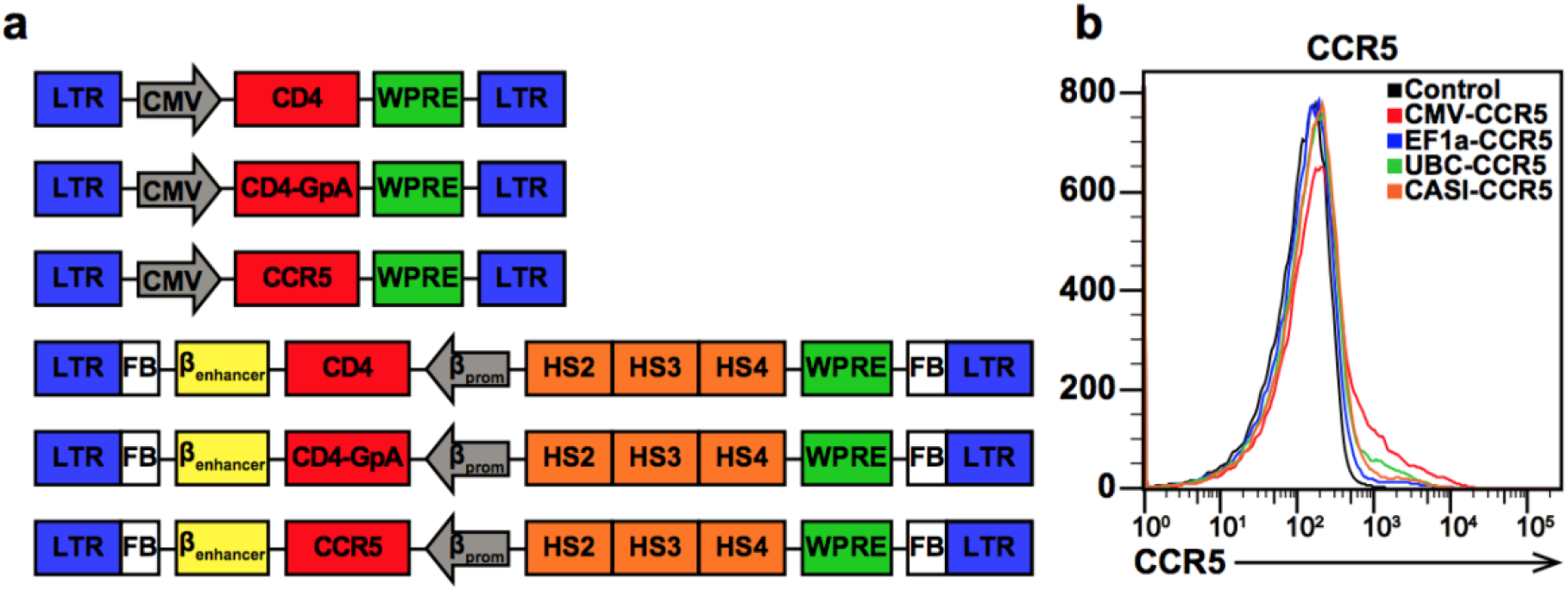
Lentiviral vector constructs for engineering RBCs. **a**, Schematic of pHAGE-based and pCCL-FB-based lentiviral vector constructs used for the delivery of CD4, CD4-GpA, and CCR5 transgenes. **b**, Comparison of ubiquitous CMV, EF1-α, UBC, and CASI promoters for the expression of CCR5 in erythroid progenitor cells on day 13 of differentiation.

**Fig. S2.**
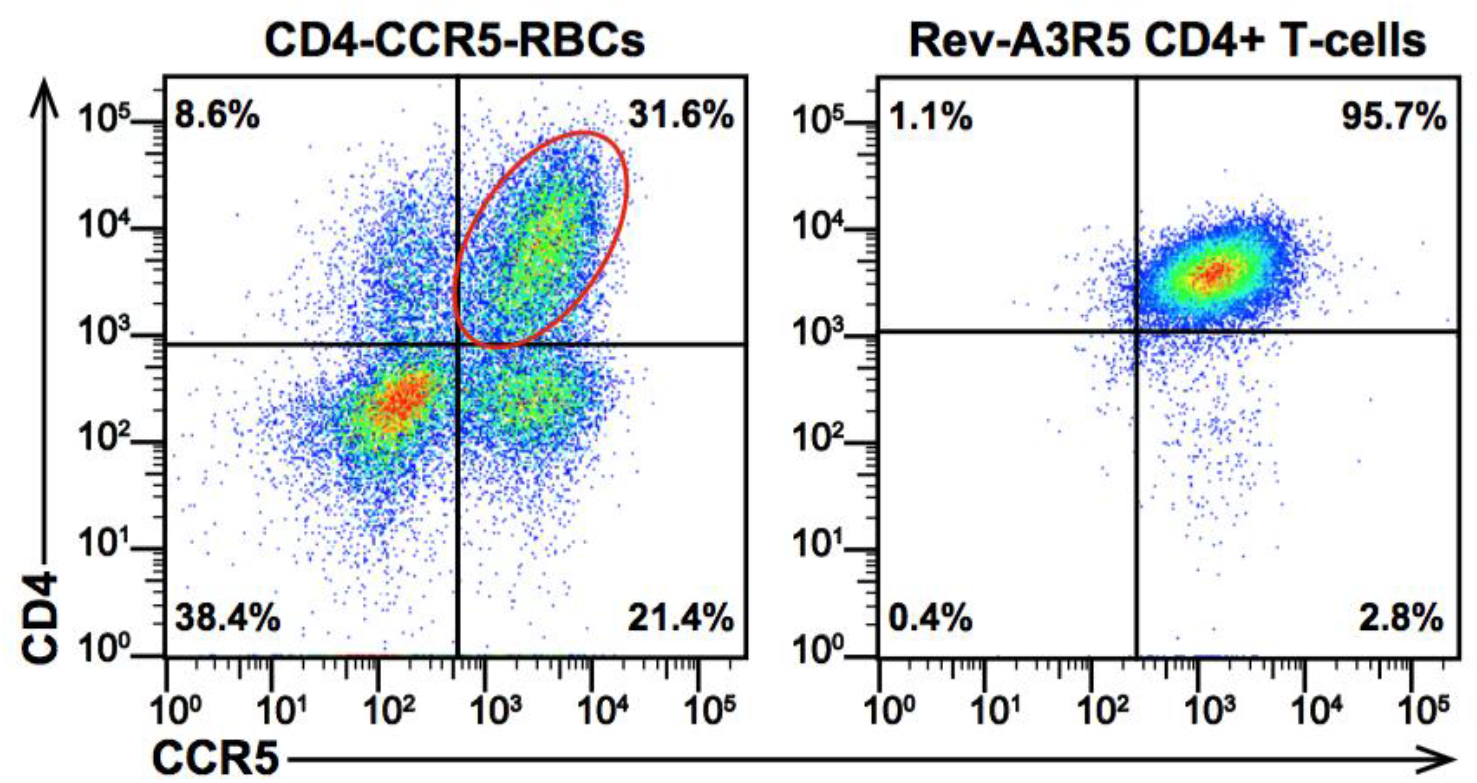
Comparison of HIV-1 receptor expression levels on CD4-CCR5-RBCs and CD4+ T-cells. Flow cytometry analysis of CD4 and CCR5 expression on enucleated CD4-CCR5-RBCs and Rev-A3R5 CD4+ T-cells.

**Fig. S3.**
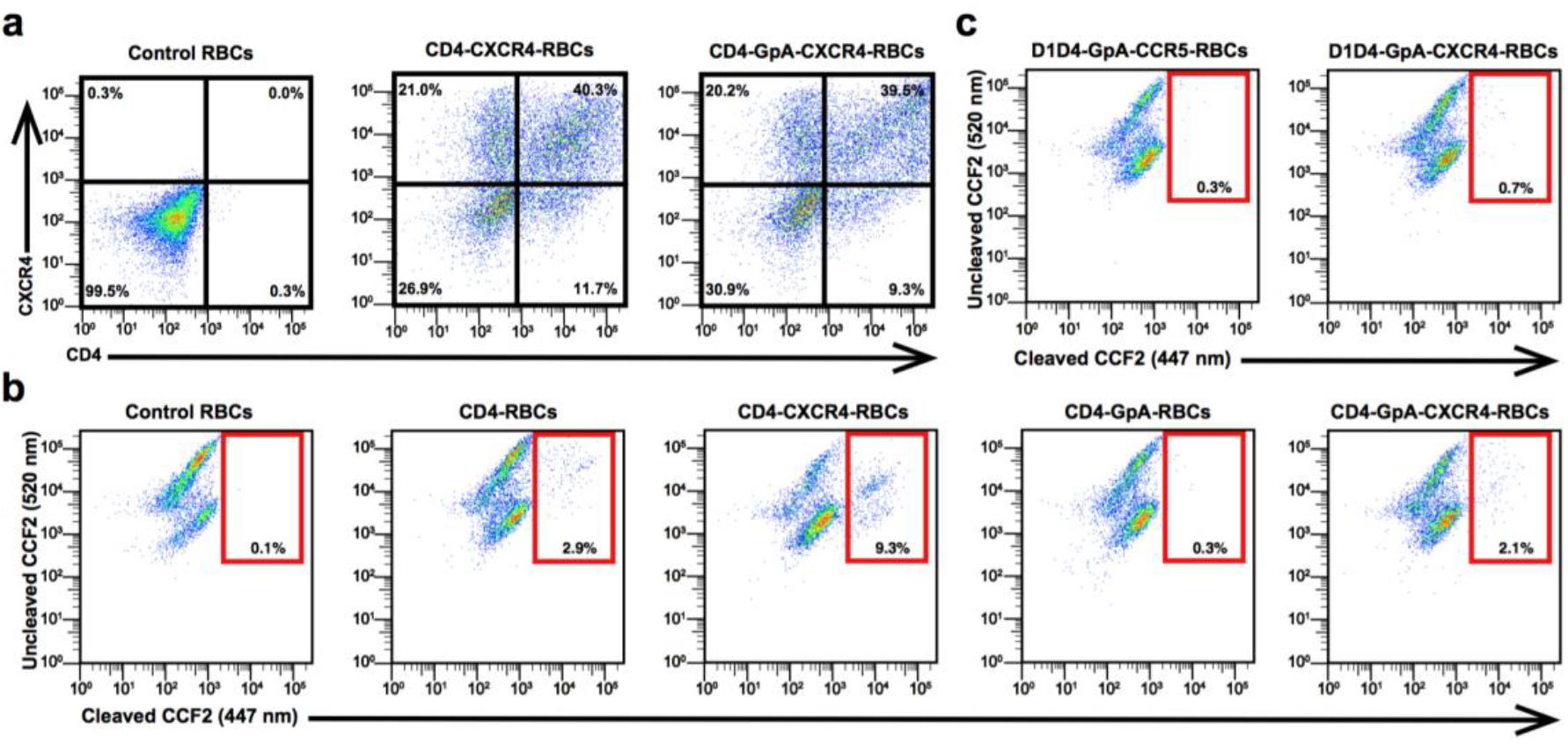
Engineered RBCs express CD4 and CXCR4 and can be infected by X4-tropic HIV-1. **a**, Flow cytometry measurement of CD4 and CXCR4 expression at the end of differentiation for control RBCs, CD4-CXCR4-RBCs, and CD4-GpA-CXCR4-RBCs. **b**, Flow cytometry analysis of HIV-1 infection of engineered RBCs after overnight incubation with an X4-tropic HIV-1_HxBc2_ pseudovirus carrying a Vpr-BlaM fusion protein. BlaM cleaves the FRET substrate CCF2-AM in infected cells resulting in a shift of its emission spectrum from green (520 nm) to blue (447 nm). **c**, Flow cytometry analysis of HIV-1 infection of RBCs expressing a chimeric D1D4-GpA fusion protein that contained the CD4 D1D4 domains to evaluate if addition of the CD4 D3D4 domains enhanced infection. BlaM assays were performed with R5-tropic HIV-1_YU2_ and X4-tropic HIV-1_HxBc2_ pseudovirus on D1D4-GpA-CCR5-RBCs (left) and D1D4-GpA-CXCR4-RBCs (right), respectively.

**Fig. S4.**
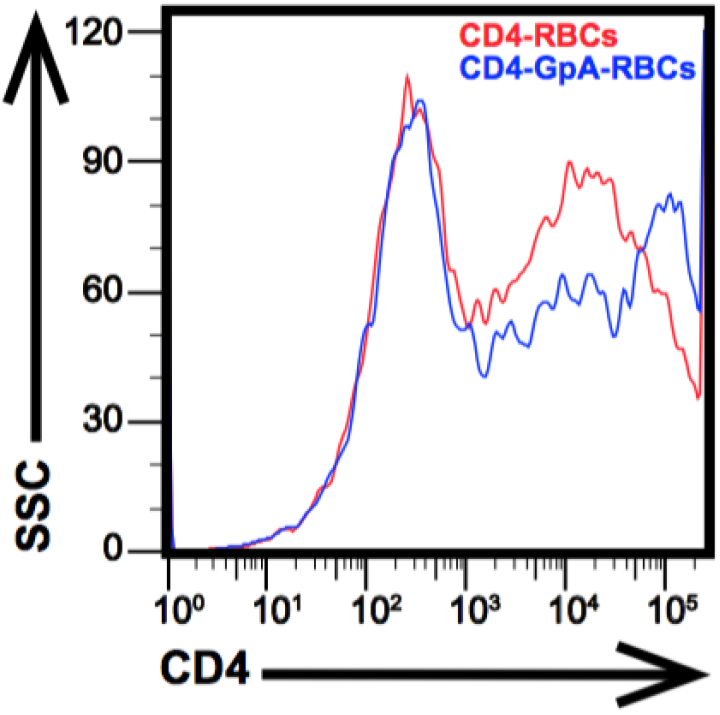
Comparison of expression levels between CD4-RBCs and CD4-GpA-RBCs. Flow cytometry analysis of CD4 and CD4-GpA expression levels on CD4-RBCs and CD4-GpA-RBCs, respectively, at the end of differentiation.

